# Development of *mPing*-based Activation Tags for Crop Insertional Mutagenesis

**DOI:** 10.1101/2020.04.22.055970

**Authors:** Alexander Johnson, Edward Mcassey, Stephanie Diaz, Jacob Reagin, Daymond R. Parrilla, Hanh Nguyen, Adrian Stec, Lauren A. L. McDaniel, Thomas E. Clemente, Robert M. Stupar, Wayne A. Parrott, C. Nathan Hancock

## Abstract

Modern plant breeding increasingly relies on genomic information to guide crop improvement. Although some genes are characterized, additional tools are needed to effectively identify and characterize genes associated with crop traits. To address this need, the *mPing* element from rice was modified to serve as an activation tag to induce expression of nearby genes. Embedding promoter sequences in *mPing* resulted in a decrease in overall transposition rate; however, this effect was negated by using a hyperactive version of *mPing* called *mmPing20*. Transgenic soybean events carrying *mPing*-based activation tags and the appropriate transposase expression cassettes showed evidence of transposition. Expression analysis of a line that contained a heritable insertion of the *mmPing20F* activation tag indicated that the activation tag induced overexpression of the nearby soybean genes. This represents a significant advance in gene discovery technology as activation tags have the potential to induce more phenotypes than the original *mPing* element, improving the overall effectiveness of the mutagenesis system.

## Introduction

Genetic improvement in crop species is facilitated by identification of genes that control important agricultural traits (Thomson et al., 2010). Using the model plant *Arabidopsis*, multiple T-DNA-based insertion mutagenesis populations have been effectively used for gene discovery in both phenotypic screens and large-scale reverse genetic approaches (Samson et al., 2002; Sessions et al., 2002; Alonso et al., 2003; Li et al., 2003). The first advantage of insertional mutagenesis over other mutagenesis techniques is that the inserted sequence, often called a tag, can be used to anchor PCR based strategies for identifying its genomic location (Parinov et al., 1999; Sessions et al., 2002). The second advantage is the ability to include sequences that interact with neighboring genes in the tag. These advanced tags contain promoter elements to induce over-expression of nearby genes [activation tags](Kakimoto, 1996; Wilson et al., 1996; Weigel et al., 2000; Dong and Von Arnim, 2003), reporter genes with splice adaptors to produce fusion proteins [gene-trap tags] (Skarnes, 1990; Sundaresan et al., 1995), or reporter genes with minimal promoters for reporting the expression patterns of nearby promoter sequences [enhancer-trap tags] (Wilson et al., 1990; Sundaresan et al., 1995). Each type of tag provides additional resources for gene function analysis and are especially important for understanding the function of essential and redundant genes that would not otherwise produce loss-of-function phenotypes. These advanced tags are critical for gene discovery in polyploid or other highly duplicated genomes including wheat and soybean.

A significant factor contributing to the success of the *Arabidopsis* insertional mutagenesis projects is the availability of inexpensive high-throughput transformation techniques (Bechtold and Bouchez, 1995; Clough and Bent, 1998). However, high-throughput T-DNA tagging is not feasible for most crop species because their transformation protocols are relatively time- and labor-intensive. Given this limitation, a number of insertional mutagenesis programs have been developed around either native or heterologous transposable elements with varying amounts of success (Aarts et al., 1995; Meissner et al., 2000; McCarty et al., 2005; Mathieu et al., 2009; Hancock et al., 2011; Cui et al., 2013). The outcomes of these programs have been hindered by the inherent transposition characteristics of the elements including preferential transposition to linked sites, low transposition rates, and tissue culture activation.

Our goal is to develop a transposon-based activation tagging system that overcomes these limitations. Activation tags designed around the *Activator (Ac)/Dissociation (Ds)* (Wilson et al., 1996; Schaffer et al., 1998; Fridborg et al., 1999; Suzuki et al., 2001; Fladung and Polak, 2012) and *Enhancer/Suppressor* (Marsch-Martínez, 2011) transposon systems engineered with constitutive promotors or enhancer sequences clearly demonstrate that modified transposons retain their mobility and also induce gene expression. These approaches have been used to clone a variety of genes including TINY (Wilson et al., 1996), LATE ELONGATED HYPOCOTYL (Schaffer et al., 1998), SHORT INTERNODES (Fridborg et al., 1999), high phenolic compound1‐1 Dominant (Schneider et al., 2005), and a strictosidine synthase (Mathieu et al., 2009). The *Ac/Ds*-based elements preferentially transpose into linked sites in many species, which requires many transformation events or labor-intensive screening measures to saturate the genome with tags (Bancroft and Dean, 1993; Nakagawa et al., 2000; Suzuki et al., 2001; Qu et al., 2008; Vollbrecht et al., 2010). In an effort to solve some of the limitations of current transposon-based activation tagging systems, we designed an activation tagging system around the *mPing* element from rice due to its favorable transposition behavior.

One of the most promising transposon-based mutagenesis systems relies on the *mPing* transposon from *Oryza sativa* (Jiang et al., 2003; Kikuchi et al., 2003; Nakazaki et al., 2003). This element has been shown to exhibit high rates of transposition in select rice cultivars (Naito et al., 2006) and can produce heritable mutations in rice (Teraishi et al., 1999; Nakazaki et al., 2003; Naito et al., 2009). When *mPing* was transferred to *Glycine max* along with the *Ping* transposase proteins required for mobilization, it produced heritable insertions without tissue culture treatment (Hancock et al., 2011). Subsequent generations of these plants revealed heritable mutant phenotypes that are being analyzed. Characterization of *mPing* transposition behavior in *Oryza sativa*, *Arabidopsis thaliana*, *Saccharomyces cerevisiae*, and *Glycine max* has shown that it transposes to unlinked sites, preferentially inserts near genes, and avoids insertion into GC-rich regions, making it an attractive candidate element to distribute activation tags across the genome (Yang et al., 2007; Naito et al., 2009; Hancock et al., 2010; Hancock et al., 2011).

Here we report the first efforts to combine the favorable transposition behavior of *mPing* with the activation tagging strategy. Before implementing such advanced *mPing*-based tags into genetically recalcitrant crop species, it is important to evaluate *mPing* for its suitability as an engineered tag. To this end, *mPing*-based activation constructs were developed and tested in both yeast and soybean. These experiments gave insight into the biology of *mPing* transposition and lead to a functional *mPing*-based activation tag platform that can be used for gene discovery in important crops.

## Results

### Development of *mPing*-based activation tags

Initially two strategies were pursued to engineer *mPing*-based activation tags.

The first approach added the ends of the *mPing* element, including the required terminal inverted repeat sequences (TIRs), to the *Agrobacterium tumefaciens Octopine synthase* (OCS) enhancer sequence. This element called *NL60*, is 100 bp smaller than the original 430-bp *mPing* element (**Figure 1a**). The second strategy was to insert multiple copies of the *Cassava vein mosaic virus* (CMV) enhancer in the center of a complete *mPing* element. This latter element (*2xE*) is 932 bp, while the version with four copies of the enhancer (*4XE*) is 1,425 bp (**Figure 1a**). To analyze the transposition frequency of these elements, a previously established yeast-based assay that has been shown to correlate with *mPing* transposition in plants was employed (Hancock et al., 2010; Payero et al., 2016). The transposition assays demonstrated that all three of these elements transposed less frequently than the native *mPing* element (**Figure 1b**). While the *NL60* element showed no transposition under these conditions, we observed a limited number of transposition events when more yeast was plated on larger plates. Together this suggests that the TIRs alone are not sufficient for robust transposition. However, the finding that the *2xE* and *4xE* activation tags also transposed at significantly lower rates suggests that other important sequences could be disrupted and/or that increasing element size reduces transposition frequency.

**Figure 1.**
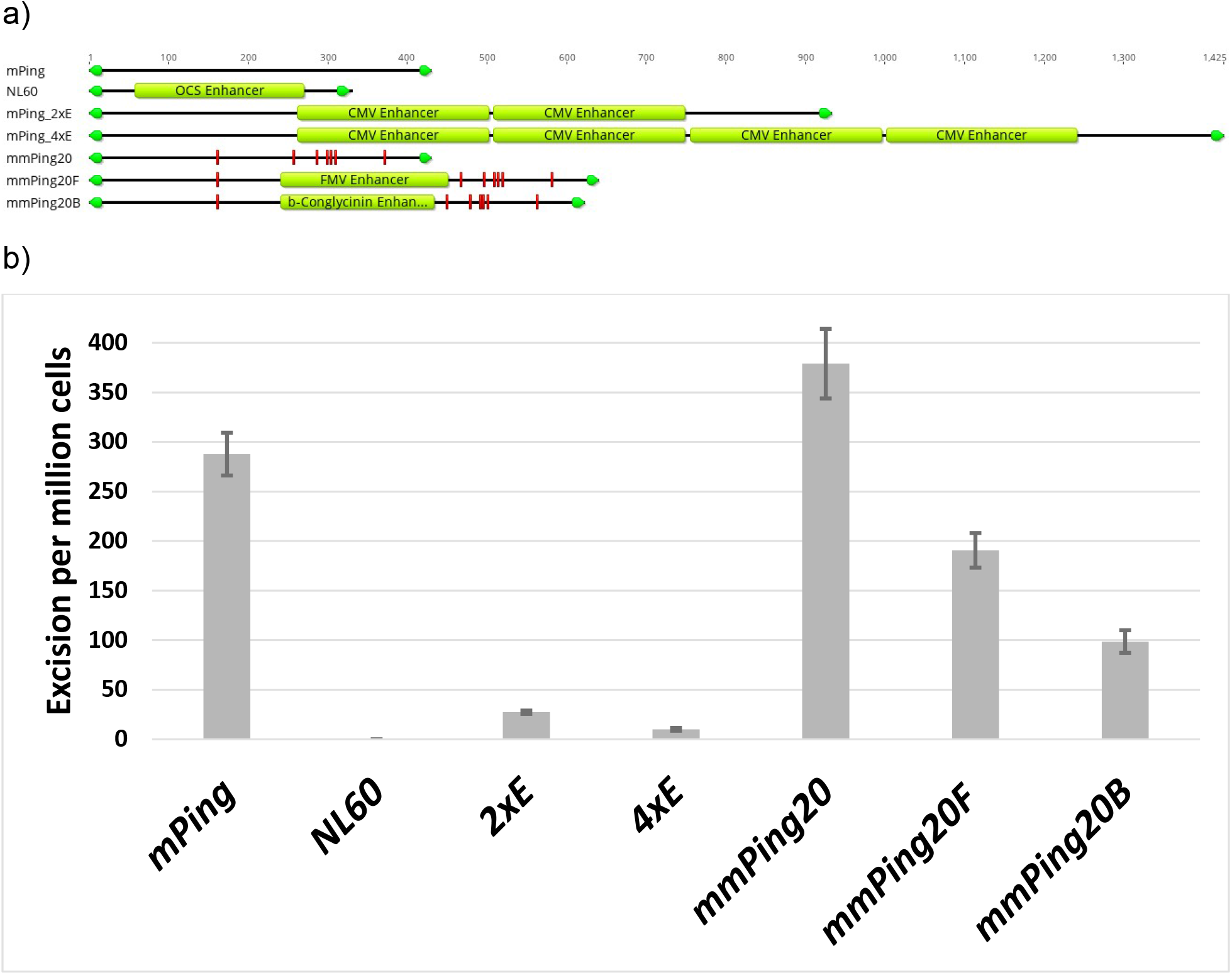
Structure of *mPing*-based activation tags and yeast transposition assays. Diagram indicating the structure of the constructs (a). TIR sequences are indicated by green arrows, red indicates mutations in the *mmPing20* element, and light green boxes indicate enhancer sequences. The images were made using Geneious version 2020.0 created by Biomatters. Yeast transposition frequency of the *mPing*, *mmPing20* and *mPing*-based activation tags (*NL60*, *2xE*, *4xE mmPing20F* and *mmPing20B*) (b). Error bars indicate the standard error of at least 6 replicates. Multiple comparisons with a Kruskal-Wallis test (nonparametric ANOVA) show that all groups differ significantly from one another (p ranging from 0.0116 to <0.0001).

### A modified *mPing* element resulted in increased transposition frequency

Transposition frequency is an important parameter in mutagenesis efficiency; therefore, mitigation of the low transposition frequency of our activation tags was addressed by minimizing their size and identifying hyperactive versions of *mPing*. Previous research showed that some native transposable elements are not optimized for transposition and genetic variants can be identified with hyperactive mobility (Yang et al., 2009). A library of 112 PCR mutated versions of *mPing,* designated *mmPing*, was screened using the yeast transposition assay. While 12 of the *mmPing* elements showed lower transposition, six were identified with higher transposition frequency. One of these, the *mmPing20* element, transposed at a significantly higher frequency than *mPing* (**Figure 1b**). This hyperactive element was found to have seven mutations relative to the original *mPing* element (T162C, T258A, T287C, T300A, T304A, T310A, G372A). The identification of the hyperactive *mmPing20* element suggests that *mPing* contains sequences that inhibit transposition complex formation.

The *mmPing20* element was used as the basis for two novel activation tags containing enhancer sequences from the promoters of the figwort mosaic virus [FMV, 207 bp] (Maiti et al., 1997) and the soybean β-conglycinin gene [182 bp] (Chen et al., 1988). The transposition of these activation tags, called *mmPing20F* and *mmPing20B* respectively (**Figure 1a**), were then analyzed using the yeast transposition assay (**Figure 1b**). The results of the assays indicate that the *mmPing20F* and *mmPing20B* activation tags transpose at a lower rate than *mPing* but at significantly higher frequencies than the *2xE* element. The finding that *mmPing20F* transposes better than *mmPing20B* even though they are similar in size (640 bp and 622 bp) suggests that the enhancer sequences can affect the efficiency of functional transposition complex formation.

### *mPing*-based activation tags can be mobilized in soybean

In order to test the mobilization of these activation tags in plants, a set of seven independent transgenic soybean lines with an *mmPing20F* activation tag construct (pEarleyGate 103 *mmPing20F*, **Figure 2a**) was developed. Transformants containing the pEarleyGate 103 *mmPing20F* construct were crossed with five transgenic lines that carry expression cassettes of ORF1 and TPase that reside in the vector pWMD23 (**Figure 2b**). A total of 20 transgenic stacks with both constructs were generated (**Figure 3a**). These were tested for transposition using PCR with primers flanking the *mmPing20F* element within the vector backbone (**Figure 2a**). Eighteen (90%) of the transgene stack combinations produced both a 960-bp and a 317-bp PCR product, indicating that somatic *mmPing20F* excision was occurring in the F1 generation (**Figure 3a**). Subsequent genotyping via PCR in the F2 generation from ten lineages that displayed transposition in the F1 generation with *mmPing20F* flanking primers was performed (**Figure 3b**). This led us to select the 16-28-3 line because it produced plants that only had the 317-bp PCR product, consistent with germinal excision of *mmPing20F* (**Figure 3b**). Progeny from line 16-28-3 were sown to identify homozygous lineages for pWMD23 and segregating for a single locus containing the pEarleyGate 103 *mmPing20F* transgenes (i.e. 16-28-3-14-4). A second line was identified that had lost the pEarleyGate 103 *mmPing20F* allele but was homozygous for the *mmPing20F* element (16-28-3-14-2). Together, this inheritance pattern indicates that a germinal transposition event occurred in the 16-28-3-14-2 line. These results, along with evidence for element mobility in transgenic soybean containing the *2xE* activation tag in two other soybean lines (**Supplemental Figures 1-3**) indicates that *mPing*-based activation tags are mobile in the soybean genome.

**Figure 2.**
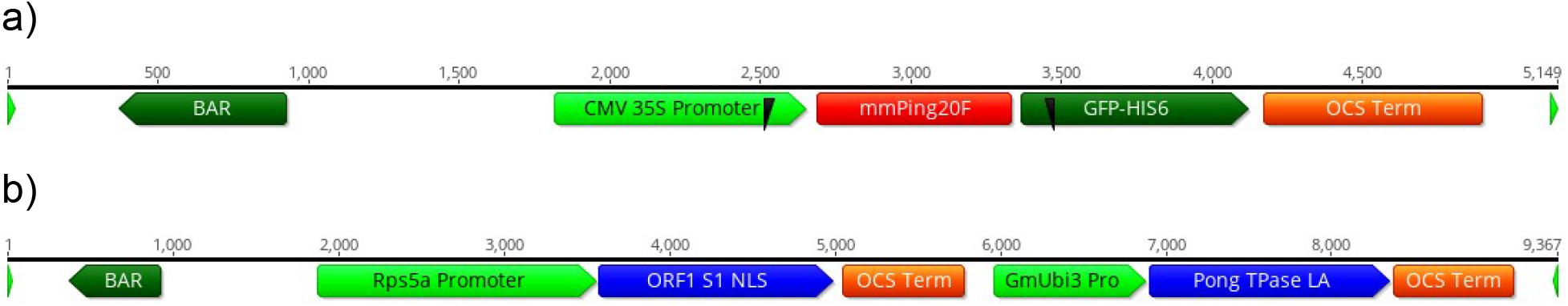
Plasmid maps. Diagrams depicting the T-DNA portions of pEarleyGate 103 *mmPing20F* (a) and pWMD23 (b). Black wedges indicate the position of the primers used to detect excision of the *mmPing20F* element. The Image was made using Geneious version 2020.0 created by Biomatters.

**Figure 3.**
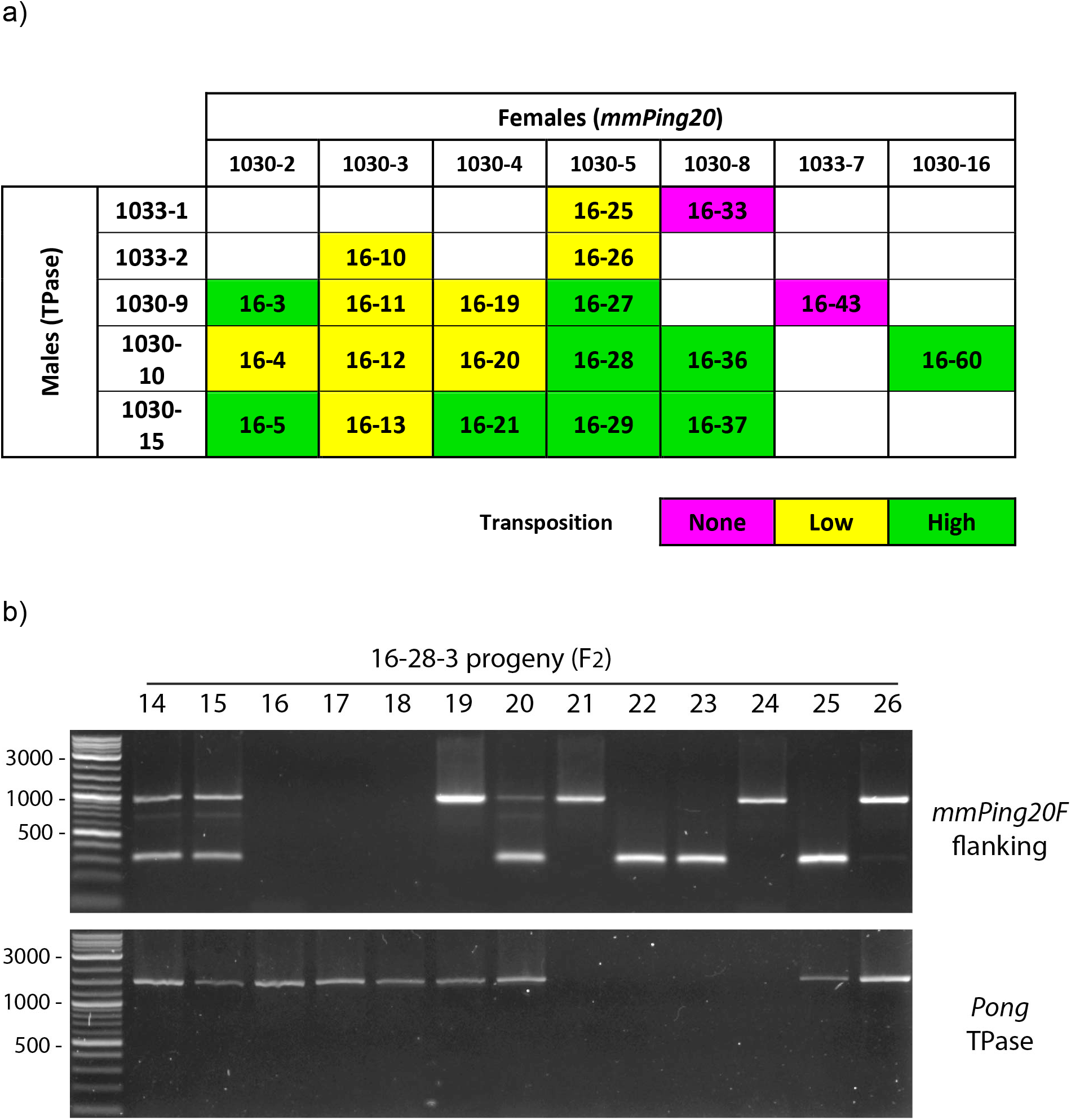
Development and analysis of the *mmPing20F* soybean population. Diagram showing the crosses that were made between the transgenic lines carrying the *mmPing20F* construct and the pWMD23 (TPase) construct (a). Colors indicate the amount of transposition detected in the F1 generation from each respective cross. PCR analysis of the 13 plants from the F2 generation of the 16-28 line (b). In the PCR with *mmPing20F* flanking primers, the upper band (960 bp) indicates that the element is still located in the transgene, while the lower band (317 bp) indicates *mmPing20F* has excised. *Pong* TPase primers indicates if the TPase expression construct pWMD23 is present.

### Insertion Site Analysis

A high-throughput sequencing approach was employed to identify *mmPing20F* insertion sites over three generations of the transgenic events. An average of 280,267 clean reads with a standard deviation of 129,024 were generated for 16 plants derived from 16-28-3 **(Supplemental Table 1)**. Variation in read depth did not likely contribute to variation in *mmPing20F* reads retrieved **(Supplemental Figure 4),** suggesting that this read depth was sufficient for the unbiased discovery of *mPing* insertions. We identified 11 *mmPing20F* insertions in this population, and each of the mappable contigs were found to be located within 4 kb of annotated genes **(Figure 4)**. Contigs that did not align to the soybean reference genome were composed of reads that matched the transformation vector, representing *mmPing20F* elements that did not mobilize. Seven of the eight plants harboring *mmPing20F* insertions had the same insertion at position 2785626 on chromosome 8 **(Figure 5)**. This insertion was 3792 bp upstream of the nearest gene, Glyma.08G035000, and 5558 bp downstream from the next closest gene, Glyma.08G035100. This insertion was present in the progeny of 16-28-3-14 and 16-28-3-15, suggesting that they were inherited through their most recent common ancestor, 16-28-3 **(Figure 5)**. However, the insertion at Chr08:2785626 was not detected in the parental lines 16-28-3-14 and 16-28-3-15 or the grandparent 16-28-3 using contig or single read alignments (**Supplemental Figure 5**). None of the other insertions were found in multiple progeny suggesting that they were either somatic insertions specific to the tissue used to generate the DNA preparation or novel germline insertions specific to that lineage (**Figure 5**).

**Figure 4.**
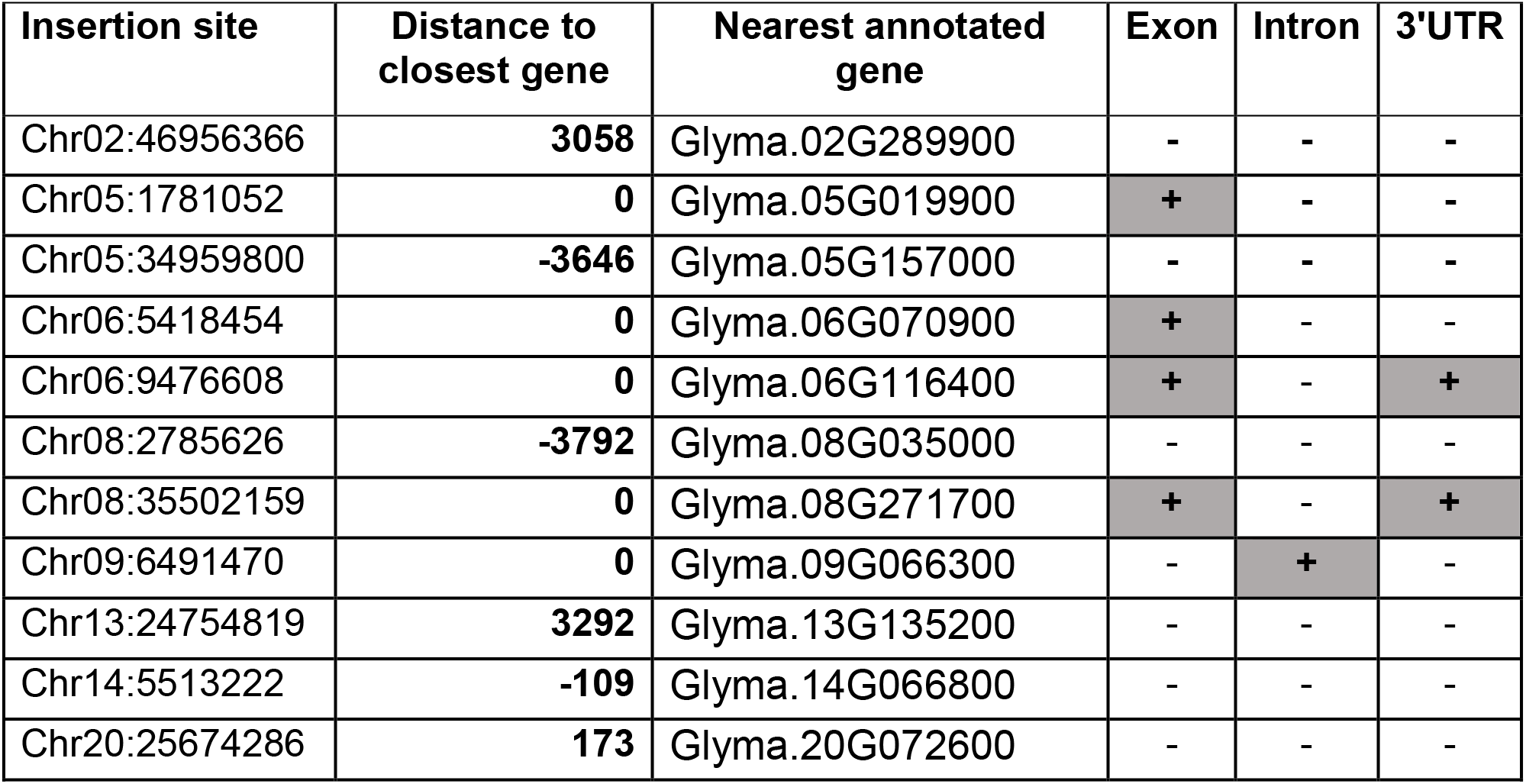
Genomic features associated with *mmPing20F* insertions identified by sequencing. The insertion site corresponds to the base immediately flanking the *mmPing20F* tag. Distances of zero indicates the insertion is within the annotated gene model. Negative distances denote upstream insertions. Shaded cells with the “+” sign indicates the insertion is in the gene feature.

**Figure 5.**
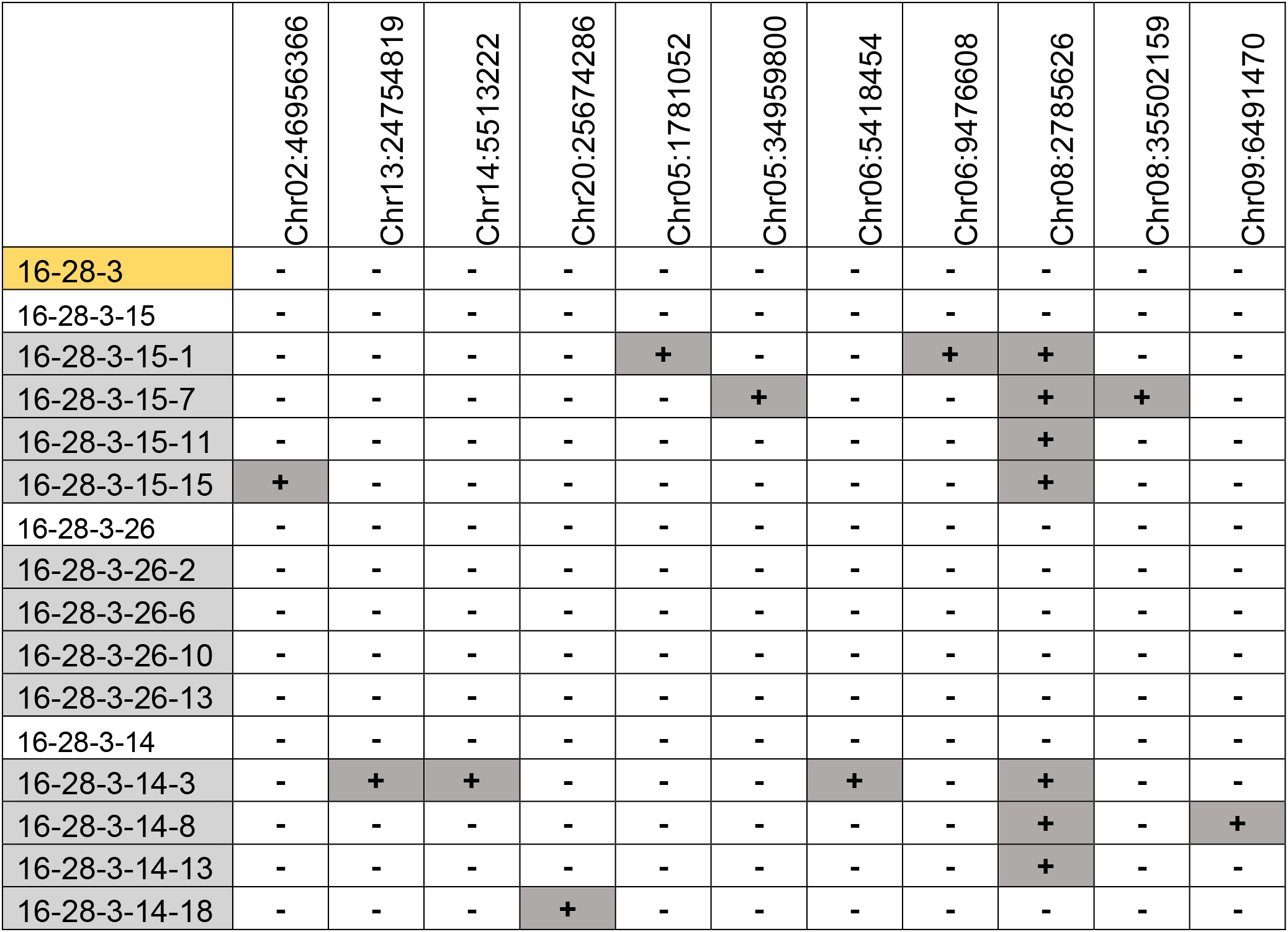
*mmPing20F* insertions detected in multiple generations. Each column indicates the absence “−“ or insertion “+” at specific loci. The original progenitor (yellow cell), three first generation progeny (white), and 12 second generation progeny (grey cells) were analyzed.

### Expression Analysis

To identify potential gene expression changes induced by the *mmPing20F* insertion, RNA-seq analysis was performed on shoot tips from progeny of two control lines (untransformed Thorne and 16-28-3-14-4 #5 [homozygous for pWMD23, null for *mmPing20F*] and two lines that have the mobilized activation tag insertion on chromosome 8 (16-28-3-14-2 #1 progeny and #2). Analysis of gene expression within a ~200-kb window surrounding the new *mmPing20F* insertion site demonstrated significant upregulation of two genes. Glyma.08G035000 (3792 bp downstream of the insertion) was upregulated 6.2 fold (padj=0.0000003), and Glyma.08G035100 (5558 bp upstream of the insertion) was induced 4.1 fold, (padj=0.02) (**Figure 6**). The remaining genes in the ~200 kb region showed no significant increase in expression. Glyma.08G035000 is annotated as an ethylene-responsive element binding protein (EREBP)-like factor, and Glyma.08G035100 is annotated as an exostosin family protein (https://phytozome.jgi.doe.gov/). Though we observed a change in gene expression for these two genes, we did not notice phenotypic changes to plant growth or architecture in the field or greenhouse. In addition to these two upregulated genes, we detected 32 additional genes unlinked to the activation tag that showed significantly different expression, including five that were upregulated and 27 that were downregulated (**Supplemental Table 2)**. These adjustments could result from the presence of the pEarleyGate 103 *mmPing20F* transgene, mutations that occurred during transformation, additional uncharacterized insertions of *mmPing20F*, or as downstream effects from the genes altered by the activation tag. GO enrichment analysis of the downregulated genes using agrigGO (http://systemsbiology.cau.edu.cn/agriGOv2/) indicated that “oxidoreductase activity” was significantly enriched (padj=0.003) **(Supplemental Figure 6)**.

**Figure 6.**
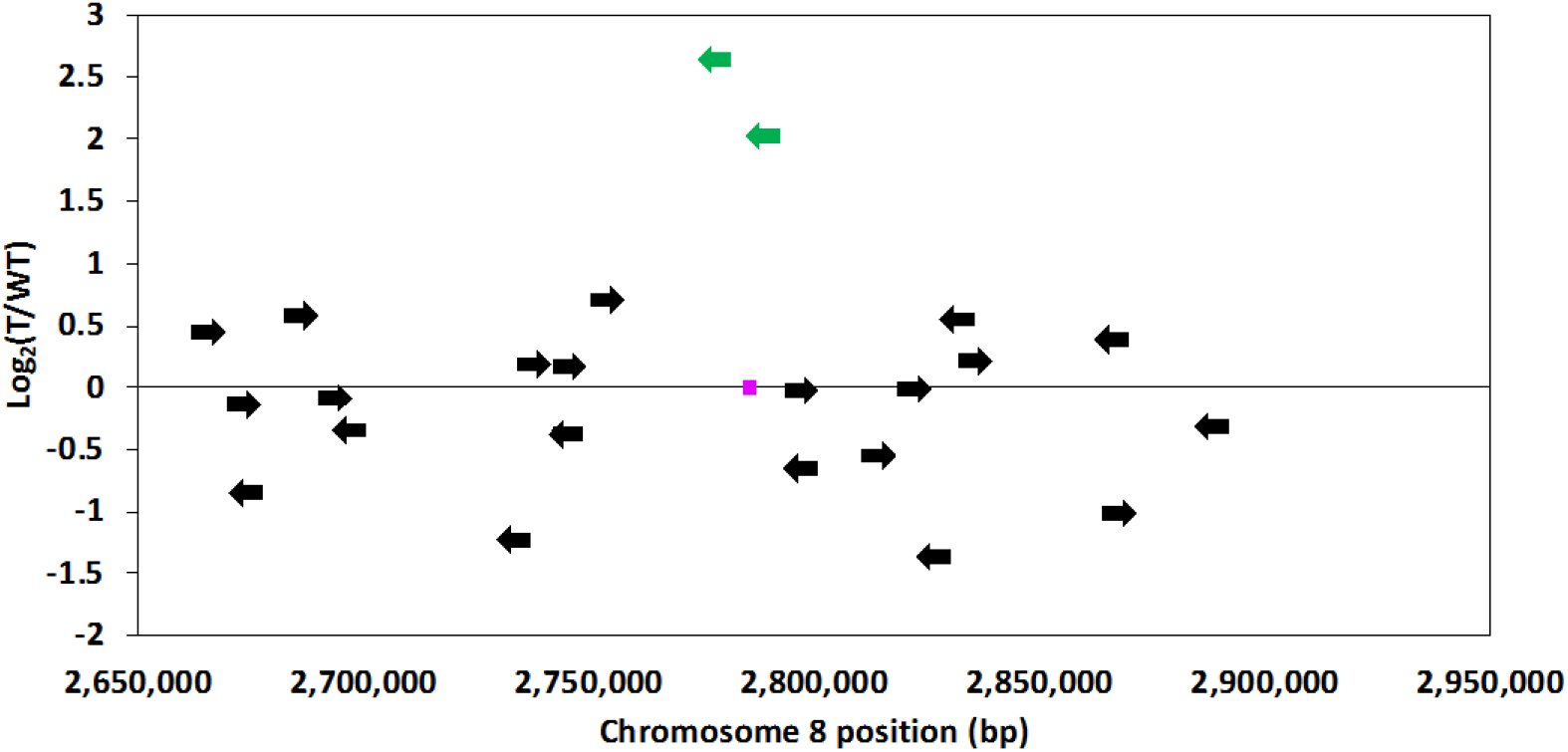
Expression changes of local genes associated with the *mPing*-based activation tag. The magenta box represents the *mmPing20F* insertion position (box not to scale). The data points correspond to annotated gene models with the arrowhead indicating transcriptional orientation (arrows are not drawn to scale). The green data points represent significantly upregulated genes (padj< 0.05) using a Wald test and Benjamini-Hochberg multiple testing correction.

## Discussion

Activation tagging mutagenesis is designed to generate miss-expression mutations. The data presented demonstrates that the *mPing*-based activation tags developed are capable of transposition (**Figure 1, Figure 3)** and altering native soybean gene expression levels (**Figure 6**). This indicates that *mPing* is a suitable vector for the delivery of enhancer sequences.

A requirement for genome-wide saturation with transposon tags is that the element must maintain mobility. Compared to other transposable elements, relatively little is known about the transposition of the *mPing* element beyond the fact that the ORF1 and TPase proteins are required (Yang et al., 2007; Hancock et al., 2010). This study provides the first report of how modification of *mPing* changes its transposition capacity. The *NL60* element, with only the 60 bp ends of *mPing*, had very low transposition rates (**Figure 1**), indicating that although the TIR sequences are sufficient for mobility, additional internal regions promote transposition activity. This finding is consistent with the results observed for a *Stowaway*-like MITE, where sub-TIR and internal regions were shown to promote transposition (Yang et al., 2009). One speculation is that these transposition promoting sequences may be involved in recruitment of the transposase proteins and subsequent formation of the transposition complex, similar to the role of *Transposon Gene A* binding sites in the transposition of *Supressor-mutator* elements (Raina et al., 1998).

The result showing that the *2xE* and *4xE* elements, consisting of a large enhancer sequence inserted into *mPing*, also had decreased transposition suggests that increases in the size of the *mPing* element results in decreased transposition rates **(Figure 1)**. This is consistent with the 430-bp *mPing* element showing considerably higher activity than the autonomous 5341-bp *Ping* element in rice (Kikuchi et al., 2003; Naito et al., 2006). This effect of size has also been observed for other Type II transposable elements (Way and Kleckner, 1985; Tosi and Beverley, 2000) and is likely one of the reasons that MITEs achieve higher copy numbers than their corresponding larger autonomous elements. Thus, the overall size of the element must be considered as additional *mPing*-based tags are constructed.

We identified both germinal and somatic insertions by introducing *mPing*-based activation tags into soybean. The majority of insertions were specific to individual plants, suggesting that they are somatic insertions that only affect a limited number of cells. However, we did identify a heritable mutation in our relatively small population that must have occurred in germinal tissue. Analysis of additional generations will be needed to calculate the frequency of novel germline insertions. This initial population suggests that germline insertion rate may be relatively low, consistent with a previous *mPing* population that showed a germline insertion frequency of about one per generation (Hancock et al., 2011). Thus, future efforts will need to focus on mechanisms to increase the transposition activity. Our goal is to increase transposition rates in soybean to those comparable to transposon tagging in maize, where populations mutagenized with an *Enhancer/Suppressor* transposon-based activation tag showed germinal excision rates ranging from 20% to 60% (Davies et al., 2019).

Interestingly, we did not detect the insertion at position 2785626 on chromosome 8 in plant 16-28-3 or its progeny, but this insertion was detected in two separate families derived from this event (progeny from 16-28-3-14 and from 16-28-3-15 contained the insertion). The fact that this insertion was not detected in 16-28-3 is not surprising, as the DNA collected for sequencing was sampled early in development, and the transposition may have occurred later and in a different tissue. It is surprising that we did not detect the insertion in the 16-28-3-14 or 16-28-3-15 plants, given that their progeny were positive for the insertion. One explanation for this is that the tissue that gave rise to the leaves sampled for sequencing in 16-28-3-14 and 16-28-3-15 had an early somatic transposition reverting the chromosome 8 locus back to wild type, but the germline insertion remained giving rise to the seeds that contained the insertion in the next generation. We did not take multiple tissue samples from these plants to test this possibility. This pattern is supported in previous transposon display analysis of native *mPing* elements where germinal bands present in the T0 generation were either absent or weaker in the T1, but the subsequent T2 plants had strong bands (Hancock et al., 2011). It is also worth noting that these plants tested positive for *ORF1* and *TPase* that are required for mobility.

Based on the behavior of the original *mPing* element, we anticipated approximately 50% of the insertions would fall within 2.5 kb of a gene (Hancock et al., 2011). We observed that seven out of the 11 mapped insertions (64%) were within 2.5 kb of an annotated gene and the remaining four were less than 3.8 kb away (**Figure 4**). This is relevant, given that our results with *mmPing20F* (FMV enhancer) showed that a gene as far away as 5.5 kb showed a significant increase in expression (**Figure 6**). We suspect that the degree of gene induction is dependent on the enhancer sequence used and its chromosomal context. Also, a larger population of *mmPing20F* mutagenized plants will need to be analyzed before we can determine the proximity limits for affecting gene expression. Nonetheless, these results are consistent with reports from T-DNA activation tagging with a cauliflower mosaic virus 35S enhancer sequence that induced overexpression at distances of up to 3.6 kb in *Arabidopsis* (Weigel et al., 2000) and 13.1 kb in tobacco (Liu et al., 2015).

The *mmPing20F* activation-tag induced upregulation of Glyma.08G03500 which is related to the *Arabidopsis TINY2*. *TINY2* belongs to the APETALA2/ethylene response factor transcription factor family. There are three TINY homologs in *Arabidopsis,* and triple knockouts grow larger than wild-type, while overexpression lines display stunted growth (Xie et al., 2019). There are two additional soybean genes with high sequence similarity to Glyma.08G035000, and phylogenetic analysis revealed four other related soybean genes cluster closer with *AtTINY1* and *AtTINY2* demonstrating that Glyma.08G035000 is only part of a larger TINY-like gene family (**Supplemental Figure 7**). Our soybean plants overexpressing Glyma.08G035000 did not display any reduction in stature or other obvious visual phenotypes under normal growing conditions. This may be attributed to subfunctionalization of the Glyma.08G035000 gene as well as the difference in magnitude of upregulation by *mmPing20F* [6.2-fold compared to the ~200-fold increase in *TINY*-overexpression *Arabidopsis* lines] (Xie et al., 2019). *AtTINY* recognizes the DRE promoter element (A/GCCGAC) and negatively regulates brassinosteroid-mediated growth through suppression of genes that respond to brassinosteroid application (Xie et al., 2019). The number of downregulated genes in *TINY-like* overexpression lines relative to wild type was smaller (27 genes) in soybean lines compared to that of *Arabidopsis* (2247 genes) (Xie et al., 2019). Additionally, we did not detect significant enrichment for DRE elements in the promoter regions of downregulated genes in our study, suggesting that Glyma.08G035000 has different target genes than *AtTINY2,* or the magnitude of Glyma.08G035000 upregulation was not sufficient to suppress its target genes.

Results from our relatively small population shows promise that *mPing*-based tags offer less up front labor to generate mutant populations than *Ac/Ds*-based tags. In most species, *Ac/Ds* transposition largely occurs in localized chromosomal regions surrounding the transgene integration site with fewer transpositions occurring on different chromosomes (Bancroft and Dean, 1993; Nakagawa et al., 2000; Vollbrecht et al., 2010). To achieve insertions on every chromosome of a species of interest, *Ac/Ds* based tags are often screened for unlinked insertions using selectable or phenotypic markers (Suzuki et al., 2001; Qu et al., 2008). This requires researchers to handle and analyze a large number of plants to identify unlinked insertions. For example, a rice *Ac/Ds* activation tagging project screened over 3000 plants produced from 37 transformation events to find insertions that reached every rice chromosome (Qu et al., 2008). The exception is the case of soybean where *Ac/Ds* based tags were not observed preferentially inserting in linked loci (Singh, 2012). However, *mPing* transposes to unlinked loci in both *Arabidopsis* and soybean (Yang et al., 2007; Hancock et al., 2011), so this system potentially offers less labor to saturate the genome with insertions across a wider range of species. Using a transformant with a single-copy vector integration, (Yang et al., 2007) found *mPing* insertions on all five *Arabidopsis* chromosomes with nine T2 individuals. Similarly in soybean, *mPing* insertions spanned 18 of the 20 soybean chromosomes by analyzing 15 plants (Hancock et al., 2011), and we identified *mmPing20F* insertions on nearly half of the soybean chromosomes in only 8 eight F3 lines.

*Ac/Ds*-based systems have an advantage over *mPing* in that it has been reported to produce more germline insertions. Novel germinal insertions were discovered at a rate of 56% (number of unique germinal insertions/total number of progeny screened) while *mPing* in soybean yielded 22% (Qu et al., 2008; Hancock et al., 2011). Since lengthening the *mPing* element reduces transposition frequency (**Figure 1**) it was not surprising that our germinal *mmPing20F* insertion discovery rate was lower than previous reports of *mPing.* As a means to ameliorate the relatively low transposition rates, screening for additional versions of the element with enhanced transposition rates is being pursued. Given that different enhancer sequences can variably affect transposition frequency (**Figure 1**), gaining insight on how these sequences contribute to transposition complex formation may also allow for the design of elements with enhanced mobility. Moreover, by incorporating a visual reporter platform that can track germinal transposition events would allow for large numbers of heritable mutations to be identified from a massive field population. Finally, efforts to develop a system to restrict ORF1 and TPase expression in the gametes are underway. This would stabilize *mPing*-based tags in the somatic tissues, allowing for simpler analysis of insertions, and strengthen its usefulness as a mutational tagging system for crop plants.

## Conclusion

Identification of the hyperactive *mmPing20* element allowed for development of activation tags suitable for plant mutagenesis. The *mmPing20F* element was shown to be heritably mobilized in soybean, resulting in overexpression of the adjacent soybean genes. Transfer of *mPing*-based activation tagging technology to additional plant species should be relatively straightforward as our plasmids should be acceptable for most dicot species and development of monocot-compatible constructs, are underway. This represents a technological breakthrough in that this *mPing*-based system will allow for production of a mutagenized populations that can facilitate gene discovery in highly duplicated or polyploid genomes.

## Materials and Methods

### Element/Plasmid Construction

The *NL60* element was made by high fidelity PCR of the *Octopine Synthase Enhancer* sequence (GenBank: AF242881) with NL60-1 For and NL60-1 Rev primers (**Supplemental Table 3**), followed by a second amplification with NL60-2 For and NL60-2 Rev (**Supplemental Table 3**). The *2xE*, *4xE*, *mmPing20F*, and *mmPing20B* elements were made by introducing restriction sites into the center of *mPing* or *mmPing20* and then cloning the enhancer sequences in by ligation. The pEarleyGate 103 *mmPing20F* construct was made by using Gateway cloning to insert the *mmPing20F* sequence into the pEarleyGate 103 plasmid (Earley et al., 2006). The pWMD23 construct was made by replacing the 35S promoter between *Xho*I and *Stu*I in pEarleyGate 100 (Earley et al., 2006) with the *Rps5a* promoter sequence by ligation and then Gateway cloning in the *ORF1 Shuffle 1 NLS* sequence. The *Pong* TPase (L418A, L420A) expression construct was then cloned into the *Pme*I site. The pWL89A *mPing*, pWL89A *mmPing20*, pWL89A *NL60*, pWL89A *2xE*, pWL89A *4xE*, pWL89A *mmPing20F*, pWL89A *mmPing20*, pWMD23 and pEarleyGate103 *mmPing20F* plasmids are available through Addgene (#140006-140007, #145787-145795).

### Yeast Assays

Transposition assays on 100 mm plates were performed using the previously described pWL89a, pAG413 ORF1 Shuffle 1 NLS, and pAG415 Pong TPase L418A, L420A plasmids in the CB101 yeast strain [*MAT*a *ade2Δ::hphMX4 his3Δ1 leu2Δ0 met15Δ0 ura3Δ0 lys2Δ::ADE2\**] (Hancock et al., 2010; Gilbert et al., 2015; Payero et al., 2016).

### Error Prone PCR

Apex Master Mix PCR reactions (50μl) were supplemented with MnCl2, MgCl2, dCTP and dTTP to conduct manganese error-prone PCR as described (Cadwell and Joyce, 1992). The *mPing* template was amplified with *mPing* TTA For and *mPing* TTA Rev primers (**Supplemental Table 3)** in two rounds of mutagenic PCR to generate a library. The library was cloned into the *ADE2* gene by cotransforming it with *Hpa*I digested pWL89a plasmid into yeast. A total of 112 clones were screened for transposition to identify the *mmPing20* hyperactive clone.

### Soybean Transformation

Stable transformation of cv ‘Thorne’ was performed using the cot-node (Zhang et al., 1999) or half-seed method (Paz et al., 2006; Curtin et al., 2011). Transformation was verified by PCR and Southern blot analysis.

### DNA extraction

DNA was extracted from young soybean leaves using an extraction buffer composed of 100 mM Tris pH 8, 20 mM EDTA, 1.42 M NaCl, 2% CTAB, 2% PVP40 and 4 mM DIECA. The leaf tissue was ground in a bead mill with 400 μl of extraction buffer and incubated at 55°C for 30 minutes. The solution was combined with 400 μl chloroform, mixed by inversion, centrifuged, and the aqueous phase was transferred to a new tube and precipitated by adding 300 μl isopropyl alcohol followed by centrifugation. The DNA pellet was washed with 70% ethanol and resuspended in TE buffer.

### Library preparation

To generate tagged libraries, 1 μg of genomic DNA was fragmented in a 20-μl reaction consisting of 2 μl of 10X fragmentase buffer (New England Biolabs) and 2 μl fragmentase (New England Biolabs) and brought up to 20 μl in nuclease free water and incubated at 37°C for 20 minutes before stopping with 5 μl of 0.5M EDTA. Fragmented DNA was purified using a Zymo clean and concentrator column (Zymo Research) following the manufacturer’s instructions and eluted into 11 μl of TLE (10 mM Tris, 0.1 mM EDTA pH 8). A 3 μl aliquot of the purified DNA fragments was blunted in a 20 μl reaction containing 2 μl buffer 2.1 (New England Biolabs), 1 μl 2 mM dNTPs, 0.2 μl T4 polymerase (New England Biolabs), and 13.8 μl nuclease-free water and incubated at 12°C for 15 min. Blunted fragments were purified using a Zymo clean and concentrator column (Zymo Research) following the manufacturer’s instructions (eluted into 8 μl of TLE) before A-tailing in a reaction containing 2 μl of GoTaq buffer (Promega), 0.2 μl 10 mM ATP, 0.8 μl GoTaq polymerase (Promega), and 7 μl of the cleaned blunted fragments (70°C for 20 min). The A-tailed fragments were size-selected using Mag-Bind® RxnPure Plus magnetic beads (Omega Bio-Tek) following the manufacturer’s instructions, except using 80% ethanol for the bead wash steps. Y-yoke adapters were prepared according to (Glenn et al., 2016) and were ligated to the a-tailed fragments “on-bead” by adding 2.5 μl of 5 μM y-yoke adapter, 2.5 μl of 10x ligation buffer (Promega), 17.5 μl nuclease-free water, and 2.5 μl of T4-ligase (Promega) [**Supplemental Table 3]** (Glenn et al., 2016). The ligation reaction was incubated at room temperature for three hours and purified by adding 25 μl of a 20% PEG 2.5 M NaCl solution, mixing, and incubation for 10 minutes at room temperature prior to magnetic separation as previously described. The ligation was eluted from the beads into 25μl TLE. Adapter ligated fragments were enriched for those containing *mPing* using a PCR primer which was reverse complementary to the *mPing* 5’ end with a tail which facilitates subsequent multiplexing and binding to the Illumina flow-cell. PCR was performed in 25 μl reactions containing 12.5 μl KAPA HiFi Hotstart ReadyMix [2X] (Kapa biosystems), 1.25 μl 5 μM *mPing* fusion primer **(Supplemental Table 3)**, 1.25 μl 5 μM iTru7 barcoded primer (Glenn et al., 2016), and 10 μl of the cleaned ligation. The PCR was performed in a thermocycler at 98°C for one minute followed by 24 cycles of 98°C for 20 seconds, 60°C for 15 seconds, 72°C for one minute, and a final extension of 5 minutes. The second index was added to the amplified libraries through a second PCR containing 12.5 μl KAPA HiFi Hotstart ReadyMix (Kapa biosystems), 1.25 μl 5 μM p7 primer **(Supplemental Table 3)**, 1.25 μl 5 μM iTru5 barcoded primer (Glenn et al., 2016), and 10 μl of the *mPing*-enrichment amplicon. The thermocycler conditions were the same as the *mPing*-enrichment PCR, except with 20 amplification cycles. The final amplified library was purified using Mag-Bind® RxnPure Plus magnetic beads (Omega Bio-Tek) following the manufacturer’s instructions, except using a bead-solution:sample ratio of 0.7:1 v:v. The cleaned libraries were quantified using the KAPA qPCR library quantification kit (Kapa biosystems), pooled in equimolar amounts, and sequenced on an Illumina Miseq with paired-end 300-bp reads.

### Bioinformatic analysis

Reads were quality-trimmed using Trimmomatic (Bolger et al., 2014) with a sliding window of 10 bp with an average quality score of 20 on the phred 33 scale. The ends of the reads were also trimmed with a quality threshold of 10. Next the reads were filtered based on whether they contained the *mPing* sequence and the target site duplication sequence TTA or TAA. The *mPing* sequence was clipped from the filtered reads using the “headcrop” function of Trimmomatic (Bolger et al., 2014), leaving the soybean sequence that flanked the insertion site. The clipped reads were assembled using CAP3 (Huang and Madan, 1999) set to a minimum read overlap of 16 bp with a 70% identity threshold to generate contigs. The contigs were used as queries against the soybean genome version Wm82.a2.v1 (https://phytozome.jgi.doe.gov) using BLASTn (Camacho et al., 2009) to identify the insertion position. The coordinates of the insertions were then cross-referenced to the coordinates of annotated gene models from soybean (https://phytozome.jgi.doe.gov) using the BEDTools function “closest” (Quinlan and Hall, 2010).

### RNA-seq

RNA was extracted from 5-cm meristem tips, including immature trifoliolates, from month-old greenhouse-grown plants. Tissue was ground in liquid nitrogen before using Trizol Reagent to purify the RNA according to the manufacturer’s recommendations. Frozen RNA samples from two plants harboring the *mmPing20F* insertion, an untransformed ‘Thorne’ plant, and a transformed ‘Thorne’ plant that only contained the pWMD23 plasmid (ORF1 and TPase expression) were shipped to Novogene for standard Illumina RNA-seq analysis. The latter two samples served as the control group. Differential expression analysis was conducted using the R package DEseq2 with default parameters (Love et al., 2014). P-values for differential expression of genes in a 200kb window surrounding the *mmPing20F* insertion were corrected for multiple testing using the Benjamini-Hochberg method (Benjamini and Hochberg, 1995). Fold changes were calculated as normalized expression values in plants harboring the insertion divided by the normalized value of the control plants. Expression differences of genes in this 200kb window surrounding the insertion were considered differentially expressed if they had a log2 (fold change) higher than one and their adjusted p-value was lower than 0.05.

### Accession Numbers

Vectors used in this article can be found under accession number 140006 and 140007 on AddGene (https://www.addgene.org/).

### Large Datasets

Sequencing data from transposon mapping and RNA-seq from this article can be found in the GenBank data libraries under BioProject PRJNA605018

## Acknowledgments

We would like to thank Gary Stacey and Minviluz Stacey (University of Missouri – Columbia) for their critical feedback on this project. We would also like to thank Theresa McManimon for performing the BertMN01 transformation and Priscilla Redd for help with statistical analysis. Funding was provided by NSF Plant Genome Research Program Grants # 0820769, 1127083, and 1444581.

## Supporting information

Supplemental Figure 1. Plasmid map of *2xE* activation tag

Supplemental Figure 2. PCR analysis of *2xE* activation tag in cultivar Thorne

Supplemental Figure 3. Southern Blot analysis of the *2xE* activation tag.

Supplemental Figure 4. Read depth analysis

Supplemental Figure 5. Insertion Positions Supported by Unassembled Reads

Supplemental Figure 6. GO enrichment analysis for downregulated genes in plants containing the heritable insertion at Chr08:2785626

Supplemental Figure 7. Phylogenetic tree of top BLAST hits of Glyma.08G035000.

Supplemental Table 1. Oligos used in this study

Supplemental Table 2. Sequencing summary

Supplemental Table 3. Insertion positions supported by unassembled reads

## Notes

https://www.addgene.org/

https://www.ncbi.nlm.nih.gov/genbank/

### Competing Interest Statement

The authors have declared no competing interest.

