## Supplemental Figures and Tables for "Development of *mPing*-based Activation Tags for Crop Insertional Mutagenesis"

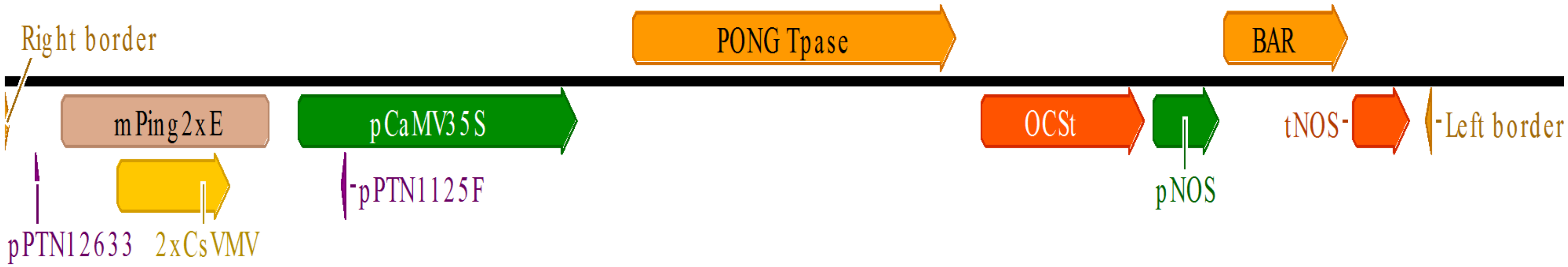

**Supplemental Figure 1**. **Plasmid map of *2xE* activation tag**

Diagram illustrating the components present in the *2xE* activation tagging T-DNA. The location of the primers used to detect excision of the *2xE* element are indicated as purple arrows labeled pPTN12633 and pPTN1125.

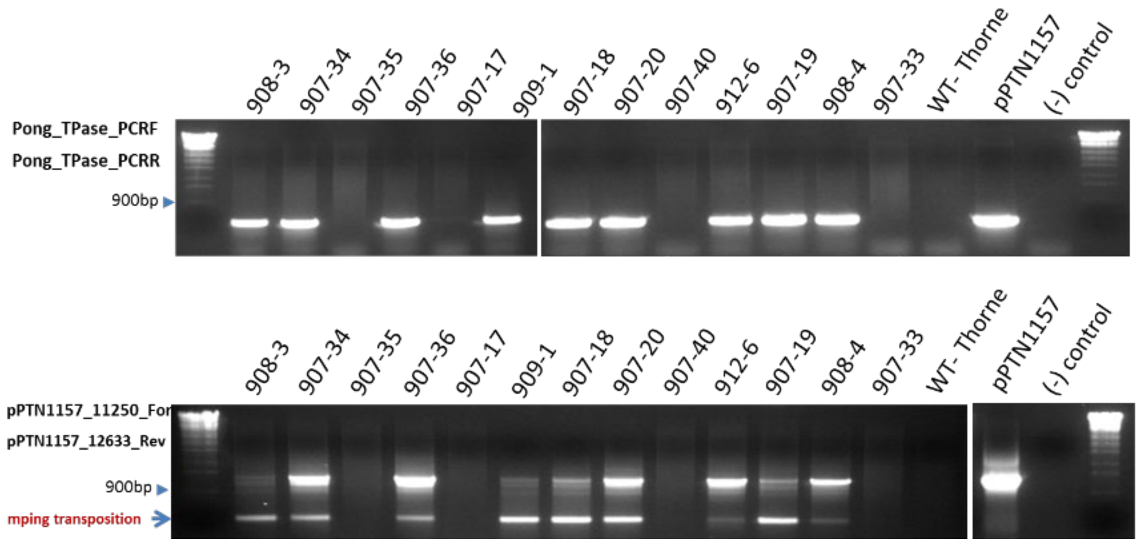

**Supplemental Figure 2. PCR analysis of *2xE* activation tag in cultivar Thorne**. PCR analysis of transgenic lines transformed with the 2xE T-DNA construct. Each lane represents an independent transgenic event. The top panel used primers that amplify a portion of the *Pong* TPase gene. The bottom panel used primers that flank the *mPing*-based *2xE* activation tag. The lower ~600bp band indicates that excision has occurred while plants that show no PCR product are escapes that do not have the transgene.

A.

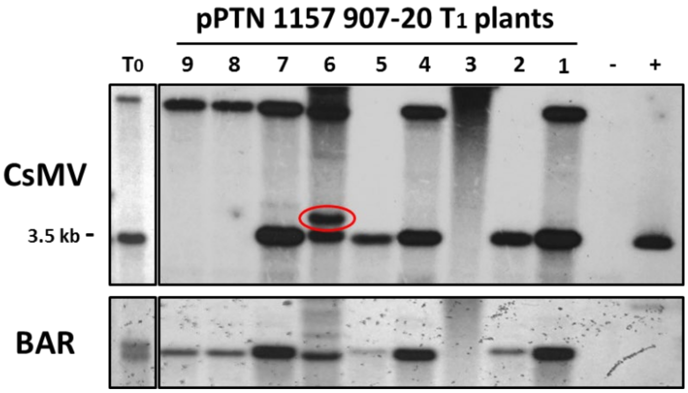

B.

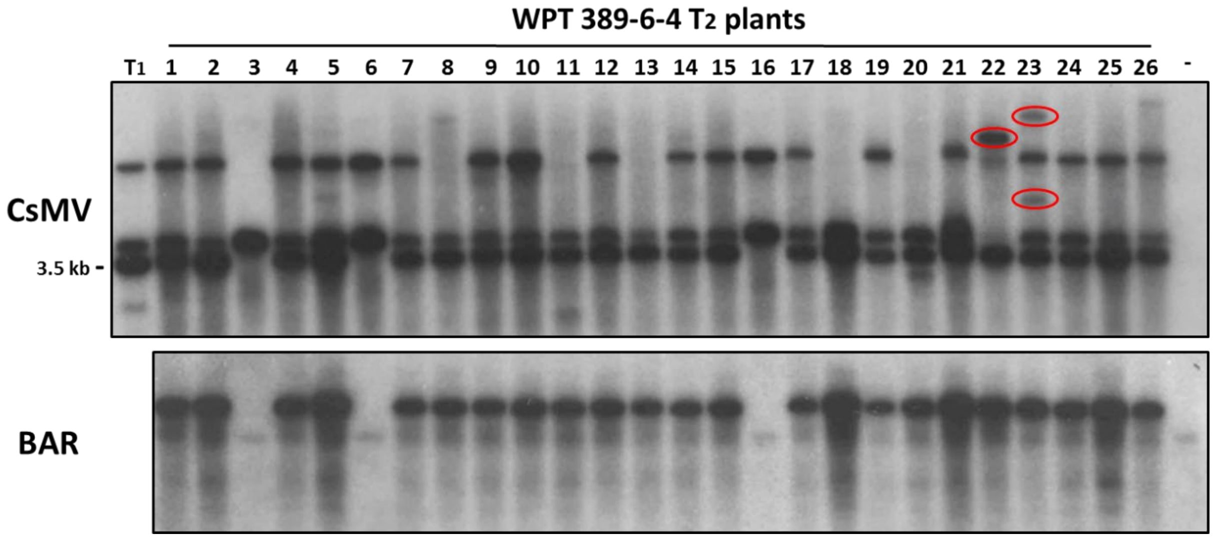

**Supplemental Figure 3. Southern Blot analysis of the *2xE* activation tag.** Autoradiograms for *Hind*III digested DNA hybridized with the CsMV enhancer (*2xE* and the vector) and BAR (selectable marker) probes. Selected transgenic lines produced in the Thorne **A)** and BertMN01 **B)** cultivars are represented. Each lane corresponds to progeny from a single self-pollinated transgenic plant. Red circles indicate novel bands not present in the parental generations (far left wells) and are evidence for *2xE* mobilization. The 3.5 kb bands indicates the *CsMV* promoter in the vector, while the other *CsMV* bands are from the *2xE* element. The negative control (-) is untransformed soybean and the positive control (+) was digested vector.

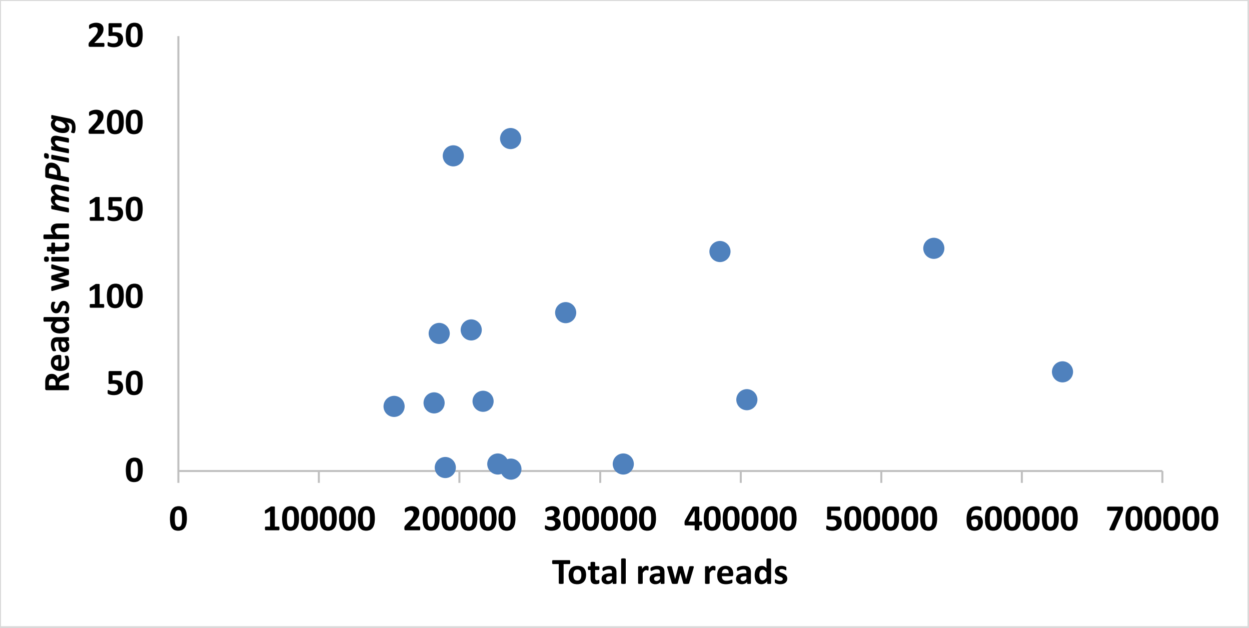

**Supplemental Figure 4. Read Depth Analysis.** The total number of reads was plotted against the total number of cleaned reads containing *mPing* and the target site duplication. R=0.1251 (Pearson coefficient) and p = 0.64

| Insertion position | 16-28-3-14-3 | 16-28-3-14-8 | 16-28-3-14-13 | 16-28-3-14-18 | 16-28-3-15-1 | 16-28-3-15-7 | 16-28-3-15-11 | 16-28-3-15-15 | 16-28-3-26 | 16-28-3-26-6 |
|---|---|---|---|---|---|---|---|---|---|---|
| Chr02:3743132 | 1 | - | - | - | - | - | - | - | - | - |
| Chr02:43335521 | - | - | 1 | - | - | - | - | - | - | - |
| Chr02:32290273 | - | - | - | - | - | 1 | - | - | - | - |
| Chr02:10633039 | - | - | - | - | - | 1 | - | - | - | - |
| Chr02:46956366 | - | - | - | - | - | - | - | 1 | - | - |
| Chr02:42419784 | - | - | - | - | - | - | - | - | 1 | - |
| Chr03:39257760 | - | - | 1 | - | - | - | - | - | - | - |
| Chr03:701314 | - | - | - | - | 1 | - | - | - | - | - |
| Chr03:7617442 | - | - | - | - | - | - | - | - | 1 | - |
| Chr04:3202029 | - | - | - | - | - | - | - | 1 | - | - |
| Chr05:1781052 | - | - | - | - | 1 | - | - | - | - | - |
| Chr05:34959800 | - | - | - | - | - | 1 | - | - | - | - |
| Chr06:5418454 | 1 | - | - | - | - | - | - | - | - | - |
| Chr06:9476610 | - | - | - | - | 1 | - | - | - | - | - |

| | | | | | | | | | | |
|---|---|---|---|---|---|---|---|---|---|---|
| Chr06:9476608 | - | - | - | - | 1 | - | - | - | - | - |
| Chr06:44472450 | - | - | - | - | - | 1 | - | - | - | - |
| Chr06:21486098 | - | - | - | - | - | 1 | - | - | - | - |
| Chr06:4875706 | - | - | - | - | - | 1 | - | - | - | - |
| Chr06:41120380 | - | - | - | - | - | - | - | 1 | - | - |
| Chr07:43972973 | 1 | - | - | - | - | - | - | - | - | - |
| Chr07:1594375 | - | - | 1 | - | - | - | - | - | - | - |
| Chr07:16643960 | - | - | - | - | - | 1 | - | - | - | - |
| Chr08:2785625 | 1 | 1 | 1 | - | 1 | 1 | 1 | 1 | - | - |
| Chr08:10162346 | 1 | - | - | - | - | - | - | - | - | - |
| Chr08:2785626 | 1 | - | 1 | - | - | 1 | 1 | - | - | - |
| Chr08:2785624 | 1 | - | - | - | 1 | 1 | - | - | - | - |
| Chr08:2786813 | 1 | - | - | - | - | - | - | - | - | - |
| Chr08:3017555 | 1 | - | - | - | - | - | - | - | - | - |
| Chr08:2785623 | 1 | - | - | - | - | 1 | - | - | - | - |
| Chr08:2785622 | 1 | - | - | - | - | - | - | - | - | - |
| Chr08:241938 | 1 | - | - | - | - | - | - | - | - | - |
| Chr08:2785616 | 1 | - | - | - | - | - | - | - | - | - |
| Chr08:2785609 | - | - | 1 | - | - | - | - | - | - | - |
| Chr08:2785619 | - | - | - | - | 1 | - | - | - | - | - |
| Chr08:2818392 | - | - | - | - | - | 1 | - | - | - | - |
| Chr08:2786480 | - | - | - | - | - | 1 | - | - | - | - |
| Chr08:2785621 | - | - | - | - | - | 1 | - | - | - | - |
| Chr08:35502159 | - | - | - | - | - | 1 | - | - | - | - |
| Chr08:713627 | - | - | - | - | - | 1 | - | - | - | - |
| Chr09:6491470 | - | 1 | - | - | - | - | - | - | - | - |
| Chr09:3906543 | - | - | - | - | - | 1 | - | - | - | - |
| Chr10:1975390 | 1 | - | - | - | - | - | - | - | - | - |
| Chr10:39003458 | - | - | - | - | - | 1 | - | - | - | - |
| Chr11:31191249 | 1 | - | - | - | - | - | - | - | - | - |
| Chr11:7662088 | 1 | - | - | - | - | - | - | - | - | - |
| Chr12:34463760 | - | 1 | - | - | - | - | - | - | - | - |
| Chr12:38687808 | - | - | - | - | - | - | - | - | - | 1 |
| Chr13:24754820 | 1 | - | - | - | - | - | - | - | - | - |
| Chr13:15400197 | 1 | - | - | - | - | - | - | - | - | - |
| Chr13:24754819 | 1 | - | - | - | - | - | - | - | - | - |
| Chr13:27807857 | 1 | - | - | - | - | - | - | - | - | - |
| Chr13:11956224 | - | 1 | - | - | - | - | - | - | - | - |
| Chr13:21552511 | - | - | 1 | - | - | - | - | - | - | - |

| | | | | | | | | | | |
|---|---|---|---|---|---|---|---|---|---|---|
| Chr13:36958813 | - | - | - | - | - | - | - | 1 | - | - |
| Chr13:27282159 | - | - | - | - | - | - | - | 1 | - | - |
| Chr14:5513222 | 1 | - | - | - | - | - | - | - | - | - |
| Chr15:8240595 | - | - | 1 | - | - | - | - | - | - | - |
| Chr16:946221 | 1 | - | - | - | - | - | - | - | - | - |
| Chr16:31433061 | - | - | 1 | - | - | - | - | - | - | - |
| Chr16:25881043 | - | - | 1 | - | - | - | - | - | - | - |
| Chr16:36269867 | - | - | 1 | - | - | - | - | - | - | - |
| Chr16:32837644 | - | - | - | - | 1 | - | - | - | - | - |
| Chr17:4438849 | - | 1 | - | - | - | - | - | - | - | - |
| Chr17:14174521 | - | - | - | - | 1 | - | - | - | - | - |
| Chr17:28991111 | - | - | - | - | 1 | - | - | - | - | - |
| Chr17:9176429 | - | - | - | - | - | 1 | - | - | - | - |
| Chr19:46442393 | - | 1 | - | - | - | - | - | - | - | - |
| Chr19:47157040 | - | - | 1 | - | - | - | - | - | - | - |
| Chr19:36692199 | - | - | 1 | - | - | - | - | - | - | - |
| Chr19:2211613 | - | - | - | - | 1 | - | - | - | - | - |
| Chr19:47504280 | - | - | - | - | - | - | - | 1 | - | - |
| Chr20:40659206 | 1 | - | - | - | - | - | - | - | - | - |
| Chr20:44943415 | - | - | 1 | - | - | - | - | - | - | - |
| Chr20:25674285 | - | - | - | 1 | - | - | - | - | - | - |
| Chr20:25674284 | - | - | - | 1 | - | - | - | - | - | - |
| Chr20:25674286 | - | - | - | 1 | - | - | - | - | - | - |
| Chr20:45866875 | - | - | - | - | - | - | 1 | - | - | - |
| Chr20:39358313 | - | - | - | - | - | - | 1 | - | - | - |

**Supplemental Figure 5. Insertion Positions Supported by Unassembled Reads.** Duplicate reads containing *mPing* sequences were collapsed into a single representative read and aligned to the soybean genome using BLAST. The presence of an insertion in one of the progeny from the top row is indicated by a “1” and absence indicated by a “-“

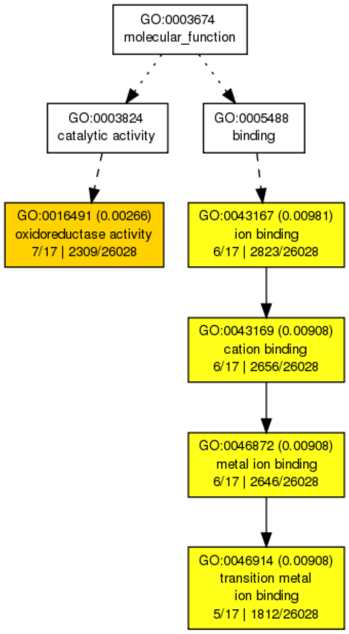

**Supplemental Figure 6. GO enrichment analysis for downregulated genes in plants containing the heritable insertion at Chr08:2785626**. The analysis was performed using agriGO (http://systemsbiology.cau.edu.cn/agriGOv2/) with the Fisher statistical method and the Hochberg multi_test adjustment method. As boxes approach red, the enrichment for the specific function becomes more significant among the test set of genes compared to background. P-values are shown in parenthesis.

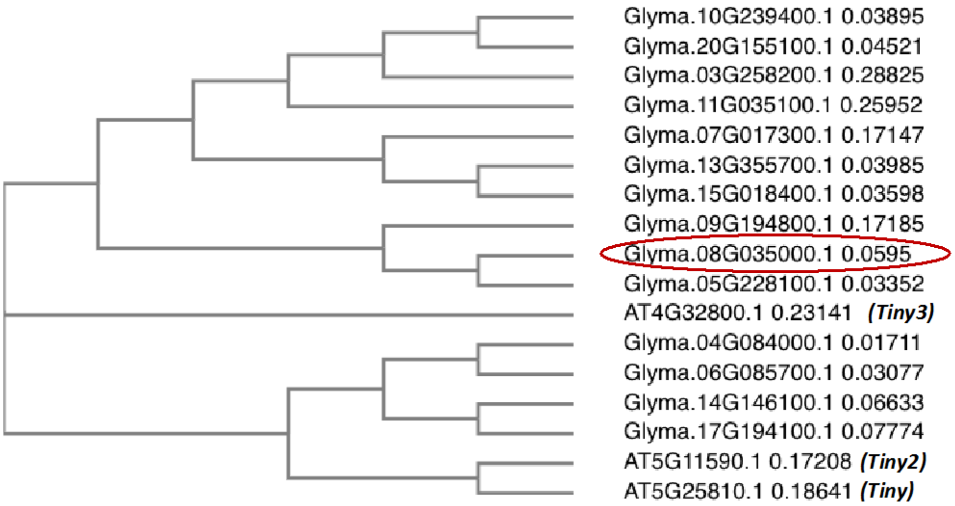

**Supplemental Figure 7. Phylogenetic tree of top BLAST hits of Glyma.08G035000.** Protein sequences were downloaded for the top 13 nucleotide BLAST hits of Glyma.08G035000.1 against the soybean genome. These protein sequences, along with the *Arabidopsis TINY*(AT5G25810.1), *TINY2* (AT5G11590.1), and *TINY3* (AT4G32800.1), were aligned by the Clustal Omega (https://www.ebi.ac.uk/Tools/msa/clustalo/) multiple sequence alignment tool to generate the phylogenetic tree. The red circle denotes Glyma.08G035000 which was the gene closest to the Chr08: 2785626 *mmping20F* insertion.

**Supplemental Table 1. Sequencing summary**

| DNA sample | total reads | total cleaned reads * | reads with mPing** | # of assembled contigs | mapped contigs*** | map %**** |
|---|---|---|---|---|---|---|
| 16-28-3 | 181,998 | 178,384 | 39 | 1 | 0 | - |
| 16-28-3-15 | 185,569 | 180,708 | 79 | 1 | 0 | - |
| 16-28-3-15-1 | 385,503 | 377,098 | 126 | 3 | 3 | 100.0 |
| 16-28-3-15-7 | 236,354 | 230,993 | 191 | 4 | 3 | 75.0 |
| 16-28-3-15-11 | 153,460 | 151,370 | 37 | 1 | 1 | 100.0 |
| 16-28-3-15-15 | 404,555 | 399,225 | 41 | 2 | 2 | 100.0 |
| 16-28-3-26 | 537,562 | 527,481 | 128 | 1 | 0 | 0.0 |
| 16-28-3-26-2 | 316,553 | 312,557 | 4 | 0 | 0 | - |
| 16-28-3-26-6 | 190,051 | 188,176 | 2 | 0 | 0 | - |
| 16-28-3-26-10 | 236,702 | 233,765 | 1 | 0 | 0 | - |
| 16-28-3-26-13 | 227,379 | 224,771 | 4 | 0 | 0 | - |
| 16-28-3-14 | 275,528 | 268,790 | 91 | 2 | 0 | - |
| 16-28-3-14-3 | 195,752 | 191,605 | 181 | 4 | 4 | 100.0 |
| 16-28-3-14-8 | 216,915 | 210,032 | 40 | 3 | 2 | 66.7 |
| 16-28-3-14-13 | 208,343 | 200,037 | 81 | 1 | 1 | 100.0 |
| 16-28-3-14-18 | 629,064 | 609,281 | 57 | 2 | 1 | 50.0 |

* reads left after adapter trimming and quality trimming
** reads that contained *mPing* sequence at the 5'end
*** number of contigs that had a high quality BLAST result from the soybean reference genome build Wm82.a2.v1
**** Percentage of contigs that returned a high quality BLAST hit

| Supplemental Table 2 Top 100 differentially expressed genes ranked by adjusted pvalue | | | | | | |
|---|---|---|---|---|---|---|
| **Gene** | **log2FoldChange** | **padj** | **Thorne**** | **T162821**** | **T162822**** | **T162845**** |
| Glyma.15G000100 | 8.40 | 2.26E-08 | 0.00 | 364.83 | 386.27 | 2.12 |
| Glyma.08G035000 | 2.63 | 2.00E-04 | 291.23 | 1533.50 | 1524.82 | 202.71 |
| Glyma.15G041100 | -9.36 | 4.86E-04 | 3808.33 | 3.37 | 2.38 | 7.43 |
| Glyma.08G182100 | -8.31 | 1.25E-03 | 1956.42 | 2.70 | 3.58 | 11.67 |
| Glyma.08G032800 | -8.91 | 1.26E-03 | 1577.12 | 2.02 | 1.19 | 7.43 |
| Glyma.20G148900 | -7.77 | 3.14E-03 | 2670.41 | 7.42 | 4.77 | 12.74 |
| Glyma.17G023000 | -8.23 | 3.14E-03 | 2341.60 | 2.02 | 5.96 | 7.43 |
| Glyma.06G266700 | -23.49 | 4.11E-03 | 568.37 | 0.00 | 0.00 | 0.00 |
| Glyma.10G159700 | -8.09 | 4.40E-03 | 5396.02 | 11.46 | 8.35 | 7.43 |
| Glyma.15G228600 | -22.82 | 6.11E-03 | 351.12 | 0.00 | 0.00 | 0.00 |
| Glyma.16G119900 | -9.43 | 6.11E-03 | 212.55 | 0.00 | 0.00 | 14.86 |
| Glyma.18G183500 | 5.48 | 7.96E-03 | 4.70 | 72.83 | 131.14 | 0.00 |
| Glyma.13G333200 | -7.48 | 7.96E-03 | 7887.93 | 34.39 | 9.54 | 22.29 |
| Glyma.12G233700 | -8.02 | 1.11E-02 | 1612.35 | 2.70 | 3.58 | 2.12 |
| Glyma.09G072000 | -7.50 | 1.11E-02 | 3740.22 | 16.86 | 3.58 | 10.61 |
| Glyma.17G069200 | -5.82 | 2.08E-02 | 13604.54 | 103.18 | 140.68 | 187.85 |
| Glyma.06G050100 | -6.23 | 2.08E-02 | 2655.15 | 5.39 | 31.00 | 45.64 |
| Glyma.11G116600 | -6.88 | 2.08E-02 | 7556.77 | 34.39 | 29.80 | 20.16 |
| Glyma.10G155600 | -7.79 | 2.08E-02 | 1344.60 | 4.72 | 1.19 | 2.12 |
| Glyma.01G051300 | -7.09 | 2.08E-02 | 2462.56 | 7.42 | 10.73 | 5.31 |
| Glyma.16G089000 | -7.03 | 2.08E-02 | 35530.34 | 79.57 | 193.14 | 74.29 |

| | | | | | | |
|---|---|---|---|---|---|---|
| Glyma.20G148100 | -7.89 | 2.19E-02 | 422.76 | 0.67 | 1.19 | 4.25 |
| Glyma.19G014400 | 1.97 | 2.63E-02 | 129.18 | 568.49 | 550.80 | 157.07 |
| Glyma.02G228100 | -6.57 | 2.82E-02 | 18226.68 | 64.74 | 127.57 | 59.43 |
| Glyma.02G119600 | -6.68 | 2.85E-02 | 623.57 | 4.72 | 1.19 | 7.43 |
| Glyma.15G096600 | -9.40 | 2.85E-02 | 219.60 | 0.00 | 0.00 | 3.18 |
| Glyma.04G041200 | -6.39 | 2.89E-02 | 1966.99 | 12.81 | 10.73 | 10.61 |
| Glyma.17G147500 | -7.41 | 2.98E-02 | 677.58 | 2.70 | 1.19 | 2.12 |
| Glyma.03G030400 | -6.74 | 2.99E-02 | 2534.19 | 14.16 | 9.54 | 6.37 |
| Glyma.18G094300 | 2.16 | 4.28E-02 | 81.03 | 504.42 | 432.77 | 128.42 |
| Glyma.07G133900 | 3.88 | 4.43E-02 | 38.75 | 223.89 | 423.23 | 5.31 |
| Glyma.03G226800 | -6.72 | 4.82E-02 | 534.32 | 2.70 | 2.38 | 3.18 |
| Glyma.13G076200 | -7.09 | 4.90E-02 | 829.07 | 4.72 | 1.19 | 2.12 |
| Glyma.07G139300 | -5.19 | 5.80E-02 | 5952.65 | 49.90 | 116.84 | 127.35 |
| Glyma.11G243400 | -5.39 | 5.80E-02 | 1325.81 | 18.88 | 13.11 | 23.35 |
| Glyma.12G106900 | -6.90 | 5.80E-02 | 430.98 | 3.37 | 0.00 | 6.37 |
| Glyma.11G214500 | -6.50 | 5.80E-02 | 4960.34 | 40.46 | 14.31 | 10.61 |
| Glyma.19G087200 | -5.79 | 5.84E-02 | 732.78 | 7.42 | 5.96 | 8.49 |
| Glyma.19G089800 | -5.09 | 5.84E-02 | 6421.20 | 57.32 | 135.91 | 147.52 |
| Glyma.15G101600 | -5.95 | 5.84E-02 | 5511.10 | 35.07 | 54.84 | 26.53 |
| Glyma.11G142900 | -5.35 | 5.84E-02 | 2265.27 | 29.00 | 27.42 | 32.90 |
| Glyma.01G112600 | 2.35 | 5.84E-02 | 79.85 | 394.50 | 681.94 | 130.54 |
| Glyma.04G030900 | -5.19 | 5.84E-02 | 1188.42 | 12.14 | 21.46 | 28.65 |
| Glyma.05G204800 | -6.75 | 5.84E-02 | 352.30 | 2.02 | 1.19 | 3.18 |
| Glyma.11G017500 | -5.51 | 5.94E-02 | 609.47 | 8.77 | 4.77 | 12.74 |

| | | | | | | |
|---|---|---|---|---|---|---|
| Glyma.15G052700 | -6.00 | 6.13E-02 | 1855.43 | 14.84 | 14.31 | 8.49 |
| Glyma.13G091100 | -4.31 | 6.14E-02 | 5309.12 | 144.31 | 138.29 | 292.91 |
| Glyma.18G141100 | -3.19 | 6.17E-02 | 1035.75 | 64.74 | 73.92 | 224.99 |
| Glyma.18G013800 | -6.33 | 6.18E-02 | 725.73 | 5.39 | 3.58 | 3.18 |
| Glyma.01G142400 | -5.02 | 6.72E-02 | 1119.13 | 25.63 | 9.54 | 35.02 |
| Glyma.11G208700 | -4.81 | 7.16E-02 | 4321.51 | 68.78 | 89.41 | 111.43 |
| Glyma.11G070600 | -5.34 | 7.16E-02 | 763.31 | 7.42 | 11.92 | 13.80 |
| Glyma.15G186100 | -5.54 | 7.16E-02 | 13967.41 | 34.39 | 271.82 | 223.93 |
| Glyma.18G052600 | -5.70 | 7.16E-02 | 1440.90 | 18.21 | 9.54 | 10.61 |
| Glyma.10G056600 | -5.93 | 7.16E-02 | 479.12 | 5.39 | 2.38 | 5.31 |
| Glyma.03G121400 | -4.83 | 7.16E-02 | 23.49 | 2.70 | 1.19 | 92.33 |
| Glyma.18G120000 | -8.51 | 7.16E-02 | 352.30 | 0.00 | 1.19 | 1.06 |
| Glyma.01G244600 | 1.69 | 7.16E-02 | 189.07 | 584.00 | 604.44 | 178.30 |
| Glyma.17G153300 | -6.60 | 7.16E-02 | 522.57 | 4.05 | 1.19 | 2.12 |
| Glyma.14G080200 | -6.29 | 7.16E-02 | 1776.75 | 15.51 | 7.15 | 4.25 |
| Glyma.15G270900 | -4.16 | 7.41E-02 | 730.43 | 23.60 | 20.27 | 56.25 |
| Glyma.08G293200 | -6.36 | 7.41E-02 | 580.12 | 1.35 | 5.96 | 3.18 |
| Glyma.12G099300 | -5.32 | 7.90E-02 | 554.28 | 8.09 | 5.96 | 10.61 |
| Glyma.09G239400 | -4.10 | 8.14E-02 | 103.34 | 4.05 | 3.58 | 28.65 |
| Glyma.04G098400 | -5.89 | 8.14E-02 | 474.43 | 3.37 | 4.77 | 4.25 |
| Glyma.07G251300 | -5.28 | 8.20E-02 | 2645.75 | 36.42 | 32.19 | 26.53 |
| Glyma.19G124600 | 2.37 | 8.93E-02 | 76.33 | 486.21 | 779.70 | 167.68 |
| Glyma.08G116900 | -2.55 | 8.96E-02 | 2261.75 | 360.11 | 182.41 | 910.58 |
| Glyma.16G057600 | -4.30 | 8.96E-02 | 5940.90 | 142.29 | 172.87 | 252.59 |

| | | | | | | |
|---|---|---|---|---|---|---|
| Glyma.19G187200 | -5.14 | 8.96E-02 | 793.84 | 12.14 | 10.73 | 13.80 |
| Glyma.03G015800 | -5.44 | 9.72E-02 | 1133.22 | 15.51 | 10.73 | 9.55 |
| Glyma.10G137200 | -5.21 | 9.72E-02 | 873.70 | 20.23 | 3.58 | 22.29 |
| Glyma.02G042500 | -8.20 | 9.75E-02 | 248.96 | 0.67 | 0.00 | 1.06 |
| Glyma.14G047500 | 1.71 | 9.88E-02 | 156.19 | 480.14 | 584.18 | 169.81 |
| Glyma.09G184600 | -4.57 | 9.91E-02 | 4829.99 | 82.95 | 125.18 | 127.35 |
| Glyma.15G244200 | -5.68 | 9.91E-02 | 577.77 | 5.39 | 5.96 | 4.25 |
| Glyma.18G005000 | -6.41 | 9.91E-02 | 799.71 | 1.35 | 8.35 | 2.12 |
| Glyma.11G000600 | 2.13 | 9.91E-02 | 159.71 | 438.33 | 650.94 | 89.15 |
| Glyma.02G136200 | -4.93 | 9.91E-02 | 702.25 | 12.81 | 10.73 | 15.92 |
| Glyma.10G017600 | -5.55 | 9.94E-02 | 10327.00 | 152.41 | 69.15 | 46.70 |
| Glyma.03G077900 | 2.33 | 1.02E-01 | 89.25 | 346.62 | 771.35 | 132.66 |
| Glyma.12G059200 | -5.47 | 1.03E-01 | 368.74 | 2.70 | 5.96 | 6.37 |
| Glyma.11G207500 | -5.11 | 1.03E-01 | 2352.17 | 42.48 | 26.23 | 25.47 |
| Glyma.20G206600 | 2.05 | 1.03E-01 | 160.88 | 1039.19 | 996.68 | 330.06 |
| Glyma.07G074400 | -5.23 | 1.03E-01 | 1930.59 | 29.00 | 22.65 | 16.98 |
| Glyma.14G177600 | 2.36 | 1.03E-01 | 153.84 | 427.54 | 1123.05 | 148.58 |
| Glyma.01G010700 | -4.84 | 1.06E-01 | 299.45 | 6.07 | 4.77 | 12.74 |
| Glyma.15G062400 | -5.37 | 1.08E-01 | 588.34 | 6.07 | 8.35 | 6.37 |
| Glyma.13G211000 | -5.43 | 1.12E-01 | 2421.46 | 12.81 | 44.11 | 16.98 |
| Glyma.05G225900 | -4.39 | 1.12E-01 | 2171.32 | 53.27 | 53.65 | 70.04 |
| Glyma.10G249500 | -5.98 | 1.13E-01 | 389.88 | 2.70 | 3.58 | 2.12 |
| Glyma.05G103300 | -4.14 | 1.14E-01 | 2282.89 | 75.53 | 59.61 | 105.07 |
| Glyma.17G258200 | -4.23 | 1.17E-01 | 5675.51 | 130.83 | 182.41 | 209.07 |

| | | | | | | |
|---|---|---|---|---|---|---|
| Glyma.17G011600 | -7.00 | 1.17E-01 | 230.17 | 0.67 | 1.19 | 1.06 |
| Glyma.12G013200 | -4.39 | 1.17E-01 | 1557.15 | 41.81 | 34.57 | 50.94 |
| Glyma.18G061100 | -3.30 | 1.18E-01 | 2185.42 | 88.34 | 166.91 | 323.69 |
| Glyma.12G112000 | -4.56 | 1.20E-01 | 2737.35 | 49.23 | 69.15 | 61.55 |
| Glyma.09G048700 | -7.10 | 1.21E-01 | 207.86 | 1.35 | 0.00 | 2.12 |
| Glyma.05G041200 | -5.19 | 1.21E-01 | 980.56 | 12.81 | 14.31 | 8.49 |
| Glyma.17G257200 | -5.04 | 1.21E-01 | 485.00 | 6.74 | 8.35 | 8.49 |

**Thorne= Untransformed control; T162821 and T162822 = Plants positive for the Chr08:2785626 *mmping20F* insertion; T162845= Transformed with pWMD23

| Supplemental Table 3. Oligos used in this study | | |
|---|---|---|
| **Name** | **Sequence 5'-3'** | **Notes** |
| iTru_A_mping_gsp2_F | ACACTCTTTCCCTACACGACGCTCTTCCGATCTGGTACCTTACCCCTATTAAATGTGCATGACACAC*C | mPing fusion primer used for PC |
| iTru_B_mping_gsp2_F | ACACTCTTTCCCTACACGACGCTCTTCCGATCTcAACACCTTACCCCTATTAAATGTGCATGACACAC*C | mPing fusion primer used for PC |
| iTru_C_mping_gsp2_F | ACACTCTTTCCCTACACGACGCTCTTCCGATCTatCGGTTCTTACCCCTATTAAATGTGCATGACACAC*C | mPing fusion primer used for PC |
| iTru_D_mping_gsp2_F | ACACTCTTTCCCTACACGACGCTCTTCCGATCTtcgGTCAACTTACCCCTATTAAATGTGCATGACACAC*C | mPing fusion primer used for PC |
| iTrusR2-stubRCp | /5Phos/GATCGGAAGAGCACACGTCTGAACTCCAGTCA*C | Component of the y-yoke adapt |
| iTrusR1-stub | ACGACGCTCTTCCGATC*T | Component of the y-yoke adapt |
| iTru_P5* | AATGATACGGCGACCACCGAGA*T | Library amplification primers (Gl |
| iTru_P7* | CAAGCAGAAGACGGCATACGAGA*T | Library amplification primers (Gl |
| iTru7_109_01 | CAAGCAGAAGACGGCATACGAGATAAGACGAGGTGACTGGAGTTCA*G | Barcoded primers (Glenn, Nilsen |
| iTru7_109_02 | CAAGCAGAAGACGGCATACGAGATACAGTTCGGTGACTGGAGTTCA*G | Barcoded primers (Glenn, Nilsen |
| iTru7_109_03 | CAAGCAGAAGACGGCATACGAGATACCGAATGGTGACTGGAGTTCA*G | Barcoded primers (Glenn, Nilsen |
| iTru7_109_04 | CAAGCAGAAGACGGCATACGAGATGTAACCGAGTGACTGGAGTTCA*G | Barcoded primers (Glenn, Nilsen |
| iTru7_109_05 | CAAGCAGAAGACGGCATACGAGATCTCGACTTGTGACTGGAGTTCA*G | Barcoded primers (Glenn, Nilsen |
| iTru7_109_06 | CAAGCAGAAGACGGCATACGAGATGACCGATAGTGACTGGAGTTCA*G | Barcoded primers (Glenn, Nilsen |
| iTru7_109_07 | CAAGCAGAAGACGGCATACGAGATGGCGAATAGTGACTGGAGTTCA*G | Barcoded primers (Glenn, Nilsen |
| iTru7_109_08 | CAAGCAGAAGACGGCATACGAGATGCCAATACGTGACTGGAGTTCA*G | Barcoded primers (Glenn, Nilsen |
| iTru7_109_09 | CAAGCAGAAGACGGCATACGAGATAAGCGACTGTGACTGGAGTTCA*G | Barcoded primers (Glenn, Nilsen |
| iTru7_109_10 | CAAGCAGAAGACGGCATACGAGATTAGTGCCAGTGACTGGAGTTCA*G | Barcoded primers (Glenn, Nilsen |
| iTru7_109_11 | CAAGCAGAAGACGGCATACGAGATTGACAACCGTGACTGGAGTTCA*G | Barcoded primers (Glenn, Nilsen |
| iTru7_109_12 | CAAGCAGAAGACGGCATACGAGATAGGAGGTTGTGACTGGAGTTCA*G | Barcoded primers (Glenn, Nilsen |
| iTru5_13_A | AATGATACGGCGACCACCGAGATCTACACGGTATAGGACACTCTTTCCCTA*C | Barcoded primers (Glenn, Nilsen |
| iTru5_13_B | AATGATACGGCGACCACCGAGATCTACACCATTGACGACACTCTTTCCCTA*C | Barcoded primers (Glenn, Nilsen |
| iTru5_13_C | AATGATACGGCGACCACCGAGATCTACACCGCAATGTACACTCTTTCCCTA*C | Barcoded primers (Glenn, Nilsen |
| iTru5_13_D | AATGATACGGCGACCACCGAGATCTACACATCCACGAACACTCTTTCCCTA*C | Barcoded primers (Glenn, Nilsen |

| | | |
|---|---|---|
| iTru5_13_E | AATGATACGGCGACCACCGAGATCTACACCTCCTGAAACACTCTTTCCCTA*C | Barcoded primers (Glenn, Nilsen |
| iTru5_13_F | AATGATACGGCGACCACCGAGATCTACACTCGATGACACACTCTTTCCCTA*C | Barcoded primers (Glenn, Nilsen |
| iTru5_13_G | AATGATACGGCGACCACCGAGATCTACACGAACCTTCACACTCTTTCCCTA*C | Barcoded primers (Glenn, Nilsen |
| iTru5_13_H | AATGATACGGCGACCACCGAGATCTACACGCTTCACAACACTCTTTCCCTA*C | Barcoded primers (Glenn, Nilsen |
| NL60-1 For | TGTGTCATGCACATTTAATAGGGGTAAGACCTGAAAGCGACGTTGGATG | |
| NL60-1 Rev | CACGGCTACCCAAAATATTATACCATCTTCTACAGGCCAAATTCGCTCTTAG | |
| NL60-2 For | TAAGGCCAGTCACAATGGGGGTTTCACTGGTGTGTCATGCACATTTAATAGGGGTAAGAC | |
| NL60-2 Rev | TTAGGCCAGTCACAATGGCTAGTGTCATTGCACGGCTACCCAAAATATTATACCATCTTC | |
| *mPing* TTA For 5’ | AGTCTCTACAATTGGGTAAGAAAACACTAAACCGTTTAGGCCAGTCACAATGGGGGTTTC | |
| *mPing* TTA Rev 5’ | ACTAAAGAATTAGCAGTCATGATTGTGAGGTCTGTAAGGCCAGTCACAATGGCTAGTGTC | |
